# Older eastern white pine trees and stands sequester carbon for many decades and maximize cumulative carbon

**DOI:** 10.1101/2020.10.27.358044

**Authors:** Robert T. Leverett, Susan A. Masino, William R. Moomaw

## Abstract

Pre-settlement New England was heavily forested, with some trees exceeding 2 m in diameter. New England’s forests have regrown since farm abandonment and represent what is arguably the most successful regional reforestation on record; the region has recently been identified as part of the “Global Safety Net.” Remnants and groves of primary “old-growth” forest demonstrate that native tree species can live for hundreds of years and continue to add to the biomass and structural and ecological complexity of forests. Forests are an essential natural climate solution for accumulating and storing atmospheric CO_2_, and some studies emphasize young, fast-growing trees and forests whereas others highlight high carbon storage and accumulation rates in old trees and intact forests. To address this question directly within New England we leveraged long-term, accurate field measurements along with volume modeling of individual trees and intact stands of eastern white pines (*Pinus strobus*) and compared our results to models developed by the U.S. Forest Service. Our major findings complement, extend, and clarify previous findings and are three-fold: 1) intact eastern white pine forests continue to sequester carbon and store high cumulative carbon above ground; 2) large trees dominate above-ground carbon storage and can sequester significant amounts of carbon for hundreds of years; 3) productive pine stands can continue to sequester high amounts of carbon for well over 150 years. Because the next decades are critical in addressing the climate crisis, and the vast majority of New England forests are less than 100 years old, and can at least double their cumulative carbon, a major implication of this work is that maintaining and accumulating maximal carbon in existing forests – proforestation - is a powerful near-term regional climate solution. Furthermore, old and old-growth forests are rare, complex and highly dynamic and biodiverse, and dedication of some forests to proforestation will also protect natural selection, ecosystem integrity and full native biodiversity long-term. In sum, strategic policies that grow and protect existing forests in New England will optimize a proven, low cost, natural climate solution for meeting climate and biodiversity goals now and in the critical coming decades.

## Introduction

A global priority for the climate has long been reducing ongoing emissions of heat-trapping greenhouse gases (GHGs) produced by burning carbon-based fuels. While essential, less attention has been given to the importance of simultaneously increasing carbon dioxide (CO_2_) removal (CDR) by natural systems. Clearing forests, draining and developing wetlands, and degrading soils account for one-third of all the CO_2_ added to the atmosphere by humans since the beginning of the industrial revolution. Together these ongoing actions continue to add approximately 1.5 PgC/year (1 Pg is 10^15^ grams or 1 billion metric tonnes). Burning wood and plant-derived liquid fuels adds even more CO_2_, and current forest management practices keep forests relatively young and limit their potential to accumulate carbon above and below ground and keep it out of the atmosphere. Two recent Intergovernmental Panel on Climate Change (IPCC) reports outlined the urgent and unprecedented imperative to halt CO_2_ emissions and remove additional CO_2_ from the atmosphere (IPCC, 2018, 2019). These reports, including the recent 1.5°C IPCC special report, identify forests as playing a major role. However, for CDR they focus primarily on afforestation (planting new forests) and reforestation (regrowing forests) and do not take into account the climate mitigation and adaptation benefits of growing existing natural forests termed “proforestation” (Moomaw et al., 2019; Cook-Patton et al., 2020).

Even achieving the goal of “zero net carbon” to limit global average temperatures to 1.5°C will still increase temperatures above the current rise of 1.1°C. This will result in additional disruption of the climate system and additional adverse consequences presently experienced; impacts are interconnected. To avoid ever-more serious consequences of a changed climate, the goal must be to become net carbon *negative* as soon as possible. Using the strategies of natural reforestation and particularly proforestation are among the most effective and least costly means for reducing the atmospheric stock of carbon as will be illustrated by the findings reported in this paper. The power of natural solutions – particularly growth and regeneration of natural forests – varies regionally, and has recently been identified as being even more robust than previously considered (Cook-Patton et al., 2020).

A second and perhaps even more urgent priority is the strong protection of intact biodiverse natural systems as outlined in the Global Assessment Report on Biodiversity and Ecosystem Services (Intergovernmental Science-Policy on Biodiversity and Ecosystem Services, 2019). This joint climate/biodiversity priority was reiterated in the peer-reviewed declaration of a Climate Emergency signed by over 13,000 scientists in late 2019 which highlighted proforestation as a global climate solution (Ripple et al., 2020), as did a recent post on behalf of the International Union for the Conservation of Nature (Kormos et al., 2020). We can use forest-based solutions to rapidly and substantially close the gap between CO_2_ emissions and removals by maximizing a range of nature-based solutions (Griscom et al., 2017), and the critical role of protecting intact ecosystems was quantified in a report that documents “wilderness” as reducing species’ extinction by half (Di Marco et al., 2019). Intact forests can simultaneously protect natural selection and biodiversity long-term, reduce extinction, and provide pathways for migration while they continue to accumulate atmospheric CO_2_ and thereby moderate temperature increases (Friedlingstein et al., 2019). Taken together, it is practical and possible to immediately protect ecosystems and prevent extinction while we increase CDR rates and accumulate additional carbon in forests and forest soils.

To date, significant attention has been focused on tropical forests (Mitchard, 2018), yet temperate forests are also biodiverse (Hilmers et al., 2018), benefit human health and well-being in highly populated areas (Karjalainen et al., 2010), and provide many essential ecosystem services (United States Forest Service, 2020a). They also have a large additional potential for CDR (Cook-Patton et al., 2020) and New England Acadian Forests are part of the “Global Safety Net” and recently identified as a Tier 1 climate stabilization area (Dinerstein et al., 2020). Current forest CDR in the United States is estimated to remove an amount of atmospheric CO_2_ equal to 11.6% of added annual CO2 equivalent emissions from the nation’s Greenhouse Gas emissions (United States Environmetal Protection Agency, 2020), with the potential for much more (Keeton et al., 2011; Moomaw et al., 2019). Consistent with the IPCC 1.5°C report that identified forests as key to increasing accumulation rates, Houghton and Nassikas estimated that the “current gross carbon sink in forests recovering from harvests and abandoned agriculture to be −4.4 PgC/year, globally” (Houghton and Nassikas, 2018). This potential carbon sink from recovering forests is nearly as large as the gap between anthropogenic emissions and removal rates, −4.9 Pg/year (Friedlingstein et al., 2019).

In the context of resource production and forest management, some carbon is stored in lasting wood products, and responsible forestry provides a reliable wood supply. However, a natural forest does not require management, and multiple analyses have found that a majority of carbon removed in a timber harvest is lost to the atmosphere. For example, Hudiburg et al. demonstrated that just 19% of the original carbon stock in Oregon forests in 1900 is in long lived wood products – approximately 16% is in landfills, and the remaining 65% is in the atmosphere as carbon dioxide (Hudiburg et al., 2019); Harmon found that the carbon storage in wood products is overestimated between 2 and 100-fold (Harmon, 2019). Furthermore, Harris et al. has shown that biogenic emissions from harvesting are 640 MtC/year, exceeding the commercial and residential building sectors, and fossil fuel emissions from harvesting add an additional 17% CO_2_ to the atmosphere (Harris et al., 2016). Strategic planning for responsible resource production can both mitigate these emissions and ensure a protected network of intact natural areas.

The US Climate Alliance highlights the importance of “net carbon accumulation” in forests across the landscape (United States Climate Alliance, 2020). This is already occurring and demonstrates the power of nature to help us restabilize the climate. A more impactful and explicit goal is to maximize carbon accumulation by utilizing some forests for responsible resource production and protecting other forests for maximal carbon accumulation for climate protection, long-term biodiversity, and human health and well-being. At a global level, if deforestation were halted, and existing secondary forests allowed to continue growing, a network of these intact forests would protect the highest number of species from extinction (Di Marco et al., 2019; World Wildlife Federation, 2020) and it is estimated that they could sequester ~120 PgC in the 84 years between 2016 and 2100 (Houghton and Nassikas, 2018). This is equivalent to about 12 years of current global fossil fuel carbon emissions, and these global numbers are conservative as outlined in recent analyses (Cook-Patton et al., 2020) and they do not factor in the enhanced regional CDR potential and high cumulative carbon that can be achieved with proforestation – for example, of carbon-dense temperate forests such as in the Pacific Northwest (Law et al., 2018) and New England (Nunery and Keeton, 2010; Keeton et al., 2011; Moomaw et al., 2019; Dinerstein et al., 2020).

Because these global and regional projections can be difficult to translate locally, particularly over time, we focused on a detailed analysis of individual trees and stands in New England. Historically, between 80% and 90% of the New England landscape was heavily forested, and early chroniclers describe pre-settlement forests with many large, mature trees reaching 1 to 1.5 m in diameter. Fast-growing riparian species like sycamores and cottonwoods could reach or exceed 2 m. Today, New England trees of this size are mostly found as isolated individuals in open areas, parks, and old estates. Old-growth forests (primary forests) and remnants and are currently less than 0.2% of northern New England’s landscape, and less than 0.03% in Southern New England, with ongoing attempts to document their value and identify their locations (Davis, 1996; Kershner and Leverett, 2004; Ruddat, 2020). Secondary forests in New England consist mostly of smaller, relatively young trees (less than 150 years, and on average less than 100 years old). Without proactive protection, and in the face of programs that almost exclusively incentivize active management (typically for young forests and/or timber production), we risk a future where the vast majority of the landscape will be managed and performing well below its carbon accumulation and biodiversity potential.

Our goal herein was to measure carbon directly in individual trees and in an “average” versus an older stand of eastern white pine (*Pinus strobus*) in New England. Most forest carbon studies focus on large geographical areas, and utilize “net” carbon data gathered from LIDAR (Light Detection And Ranging) and satellite technology, as well as statistical modeling based on the Forest Inventory and Analysis (United States Forest Service, 2020b) and Carbon On Line Estimator (National Council for Air Stream Improvement, 2020), products of the US Forest Service. We explore these options and note that carbon estimates from different tools and models can lead to disparate results at the level of individual trees – errors that can therefore be extrapolated to stands (Leverett et al., 2020). Therefore, we capitalized on the extensive tree-measuring protocols and experience of the Native Tree Society (NTS) to conduct highly accurate direct field measurements and measure volume precisely in younger vs. older trees growing in stands (Native Tree Society, 2020). We used direct measurements to evaluate volume-biomass models from multiple sources and developed a hybrid – termed FIA-COLE – to capitalize on the strengths of each model.

For all aspects of this analysis we calculated the live above-ground carbon (in tonnes) in eastern white pines and individuals of other species in a pine stand using conservative assumptions and direct measurements wherever possible and well as direct measurements of individual dominant pines up to 190 years in age. Our basic analyses likely apply to other northeastern conifers such as red pines (*Pinus resimosa*), eastern hemlock (*Tsuga canadensis*), and red spruce (*Picea rubens*).

## 2. Materials and methods

This paper centers on the study of individual eastern white pines of a representative older stand in Western Massachusetts, collectively named the *Trees of Peace (TOP)* located in Mohawk Trail State Forest, Charlemont, MA. The *TOP* has 76 pines covering 0.6 to 0.7 ha. We also collected and analyzed data from NTS measurements in 38 other sites in the East (Supplement 1). Since 1990, NTS has taken thousands of on-site direct measurements of individual trees in multiple stands of eastern white pines (*Pinus strobus*) (See Supplement 1 for list of sites). Measurements are published on the society’s website (NativeTreeSociety), comprehensive measurement protocols (Leverett et al., 2020) were adopted from those developed by NTS (Leverett et al., 2020) and incorporated into the American Forests Tree Measuring Guidelines Handbook (Leverett and Bertolette, 2014). A brief description of the measurement methods and models is provided in section 2.1, Supplement 2 and (Leverett et al., 2020).

In the pine stands, a point-centered plot was established with a radius of 35.89 m, covering 0.403 hectares (subsequently referred to as 0.4 ha), with the goal of evaluating a standard acre (radius: 117.75 ft), and thus relevant to forestry conventions in the U.S. Within the *TOP*, 44 mature white pine stems were tallied along with 20 hardwoods and eastern hemlocks down to a diameter of 10 cm at breast height. The measured acre had 50 pines in July 1989, and since then six trees were lost in a wind event. The pines are ~160 years old, and the hardwoods and hemlocks are estimated to be between 80 and 100 years old.

### 2.1 Height and diameter direct measurement methodology

We quantified the volume of the trunk and limbs of each tree from heights and diameters measured by state-of-the-art laser-based hypsometers, monoculars with range-finding reticles, traditional diameter tapes, and calipers (Leverett et al., 2020). Each high-performance instrument was calibrated and independently tested for accuracy over a wide range of distances and conditions. Absolute accuracies of the two main infrared lasers were verified as +/− 2.5 cm for distance, surpassing the manufacturer’s stated accuracy of +/− 4.0 cm. The tilt sensors were accurate to +/− 0.1°, meeting the manufacturer’s stated accuracy. The combination of these distance and angle error ranges, along with the best measurement methodology, gave us height accuracies to within 10 to 15 cm on the most distant targets being measured and approximately half that on the closest targets. We distinguished the rated and/or tested accuracy of a particular sensor of an instrument (such as an infrared laser or tilt sensor) from the results of a measurement that utilized multiple sensors.

Tree heights were measured directly for each pine with a visible top, using the sine method (Supplement 2) whenever possible rather than the traditional tangent method. Our preference for the sine method is supported by NTS, the US Forest Service (Bragg et al., 2011) and American Forests (Leverett and Bertolette, 2014). The more traditional tangent method often over/under-estimates heights by treating the sprig being measured (interpreted as the top), as if it were located vertically over the end of the baseline. The heights of 38 white pines in the *TOP* with visible tops were measured directly using the sine method.

### 2.2 Use of a form factor and FIA-COLE in determining pine volume

To compute directly the trunk volume from base to absolute top of a tree, diameters at base and breast height were measured with conventional calibrated tapes according to the procedures established and published by NTS. Diameters aloft were measured with the combination of laser range-finders and high performance monoculars with range-finding reticles. A miniature surveying device, the LTI Trupoint 300, was also used. Its Class II, phase-based laser is rated at an accuracy of +/− 1.0 mm to clear targets. In the *TOP*, we computed the volume of each pine’s trunk and limbs using diameter at breast height, full tree height, trunk form, and limb factors. (See Supplement 3 for a discussion on the development of the form factor and its importance in measuring volume, with comparisons to other methods of measurement).

Detailed measurements of 39 sample trees established an average form factor (see NTS measurements in Supplement 3, Table S3.2). The volume of each sample tree was determined by dividing the trunk into adjacent sections, with the length of each section guided by observed changes in trunk taper and/or visibility. Each section was modeled as the frustum of a regular geometric solid (neiloid, cone, and paraboloid; see Supplement 3 and Leverett et al., 2020, for formulas). Section volumes were added to obtain trunk volume, the form factor was determined needed to equal the trunk volume, given the total height and breast-high diameter of each pine. This produced an average factor that would fit the pines growing in a stand. We applied the average form factor to all pines included in the *TOP* as one determination of trunk volume.

For comparison to our direct volume measurements, we applied a hybrid volume-biomass model to compute trunk volumes for pines in the *TOP*. This hybrid allowed us to make use of the extensive analysis of the US Forest Service Forest Inventory and Analysis (FIA) program and database (which determines volume and biomass through the use of allometric equations) as well as the Carbon On-Line Estimator (COLE). This hybrid was termed FIA-COLE. See Supplement 4 for a full explanation of the variables and equations for defining trunk volume. We finalized volumes for the pines in the *TOP* by averaging our direct measurements with those of FIA-COLE.

For the total volume of the above-ground portion of a pine, we derived a factor for limbs, branches, and twigs as a proportion of the trunk volume using the FIA-COLE model (Supplement 5). That model includes all the branching in what is defined as the “top” in a biomass calculation and the limb factor for large trees is typically an additional 15-16%. We ran the model for each of the individuals in the *TOP* and calculated the volume. This was converted to biomass (density) and then to carbon mass using the carbon mass fractional factor.

### 2.3 Analysis of individual pine trees and a representative stand

In addition to the *TOP*, and older exemplary pines, we quantified above-ground carbon in younger trees and a representative stand. To determine an “average” pine at 50 years we defined two populations: (1) trees at 50 years that are still alive today, and (2) trees that were alive at 50 years, but are missing today. This allowed us to compute an average trunk size for the missing trees and the associated carbon. We also measured white pines from young to older ages to estimate growth rates and volumes.

We extensively studied an ~80-year-old stand of pines adjacent to the *TOP* (Supplement 6) growing on a terrace located just downslope from the *TOP* in an area fairly well protected from wind and with adequate soil depth. This age is more representative of the average stand of eastern white pine in New England. We also considered the range of pines of known ages from stands within the vicinity and elsewhere. Where we could, we examined ring growth and height patterns for individual pines during their early years on a variety of sites in different geographical locations. In some cases, we examined stumps and measured the average ring width. In other cases, we measured trees and counted limb whorls to get age estimates.

We measured the tallest pine in great detail and over a long time-span (referred to as Pine #58, its research tag number). Pine #58 has been measured carefully and regularly over a period of 28 years. In 1992 the tree was 47.24 m tall and 2.93 m in circumference. Since then, it has been climbed 4 times, tape-drop-measured, and volume-determined. Pine #58 continues to grow and enabled us to quantify the changes in carbon accumulation in a dominant tree over decades. See Supplement 7 for a detailed measurement history of Pine #58. Additional trees were measured at sites listed in Supplement 1. As noted, above-ground volumes were converted to mass using standard wood density tables (United States Department of Agriculture, 2009). The air-dried density for white pine is 385.3 kg/m^3^ (0.3853 tonnes/m^3^). We calculated the amount of carbon in each pine using a conservative figure of 48% of total air-dried weight (50% is used more commonly; the percentage of carbon content in different species ranges from ~48% to 52+%). Therefore we calculated a cubic meter of white pine trunk or limbs as holding 0.18494 tonnes of carbon. Note that the carbon in a cubic meter of wood varies depending on the species and is usually higher in hardwoods (United States Department of Agriculture, 2009).

## 3. Results

Using conservative assumptions where needed, and direct measurements wherever possible, we found that individual eastern white pines accumulate significant above-ground volume/carbon at least up to 190 years, that this volume/carbon accumulation can accelerate overtime, and that a stand of pines can double its above-ground carbon between ~80 and 160 years.

### 3.1 Analysis of dominant individuals and averages for stand-grown pines

As Pine #58 is the tallest and the largest tree (volume) in the *TOP,* its performance over time was analyzed in great detail. It started growing as part of a more tightly packed stand, but presently has ample space. Its circumference at breast height is 3.30 m, its height is 53.64 m, and its crown spread is approximately 15.5 m. Over a period of 26 years, beginning in 1992, Pine #58 has grown in circumference at an average rate of 1.39 cm per year and grown in height 23.71 cm per year. For a chronosequence, we assumed that Pine #58 grew a lot when it was young – an average of up to 61 cm per year in its first 50 years. Its trunk and limb volume was 23.02 m^3^ at the end of the 2018 growing season (Supplement 7).

Figure 1 shows the increase in height, circumference and volume of Pine #58 within each 50-year interval up to 150 years. Its estimated age is ~160 years, and we used a chronosequence to determine previous epochs. For dominant pines in stands on good sites, ring widths for the first 50 years average ~0.6 cm and thus a 1.88 m circumference at 50 years. (We measured one exceptional pine at 2.13 m in circumference.) Heights of stands at age 50 depend largely on the site index (the average height of a stand at 50 years), and indices for white pine on good sites usually range from 25.0 to 33.5 m. We used an index near the upper range (30.5 m) and well above the average for Massachusetts to assume rapid early growth. Based on these principles, the change in circumference and growth in height were greatest in the first 50 years, and decreased in the next two 50-year periods, confirming young pines “grow more rapidly” in terms of annual height and radial increases. However, volume growth, and thus carbon accumulation, increased with age. This is primarily because volume increases linearly with height but increases as the square of the diameter (see Figure 1 and Supplement 8).

**Figure 1.**
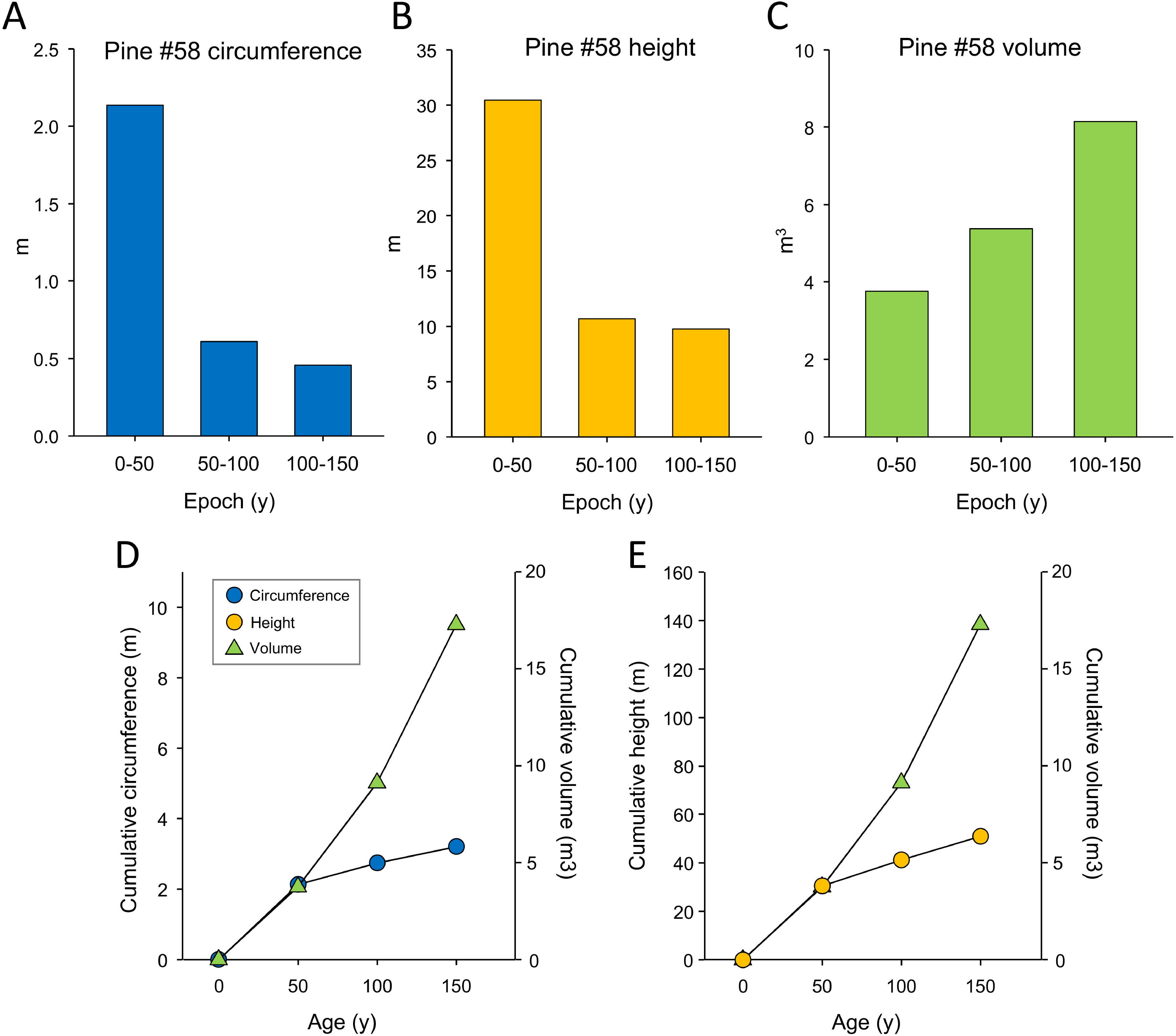
Changes in circumference, height and volume of a stand-grown individual eastern white pine (Pine #58) in three 50-y intervals. *Upper panels* - **A**: Change in circumference during 0-50, 50-100 and 100-150 y. **B:** Change in height between 0-50, 50-100 and 100-150 y. **C**: Change in above-ground tree volume (trunk plus limbs) between 0-50, 50-100 and 100-150 y. *Lower panels* - **D:** Cumulative circumference at 50, 100 and 150 y compared to cumulative above-ground volume. **E.** Cumulative height at 50, 100 and 150 y compared to cumulative above-ground volume. On each panel initial slopes were matched to reflect the rapid change in circumference and height during the first 50-y interval. Note that volume is a proxy for above-ground carbon. Values for circumference, height and volume of Pine #58 were determined by a combination of direct measurement and chronosequence and described in the text and in Supplement 7.

As noted, we assumed Pine #58 had optimal rapid growth in the first 50 years. Even so, our analysis supports the conclusion that the pine accumulated the majority of its current carbon *after age 50* and at a slightly increased rate. Pine #58 now stores 4.24 tC above ground and continues to grow. For comparison, the carbon sequestered in the highest volume 50-year-old pine that we encountered (2.13 m circumference, 34.75 m height, and 0.4353 form factor) is 1.01 tC. Therefore, even in the best-case scenario Pine #58 would have acquired less than a quarter of its current carbon by age 50.

The carbon advantage gained by the older trees accelerates with their increasing age and size, a finding that has been affirmed globally (Stephenson et al., 2014). Figure 2 documents the average volume in individual pines in the stands at ~80 and ~160 years as well as several additional large pines. MSF Pine #1, the largest pine in Monroe State Forest, western Massachusetts, has a trunk volume of 35.9 m^3^ at approximately 190 years (6.15 tC; Figure 2). Assuming its early years accumulated 1.01 tC at 50 years, which is the fastest growing 50-year old pine we measured in all sampled locations, the large pine added 5.14 tC between 50 and 190 years, or 1.84 tC per 50-year cycle after year 50. This is at least 1.82 times the rate of growth for the first 50 years. This compares to a 1.6 ratio for Pine #58. In both cases more than 75% of the carbon they sequestered occurred *after* their first 50 years even when assuming the most optimal growth observed during the first 50 years.

**Figure 2.**
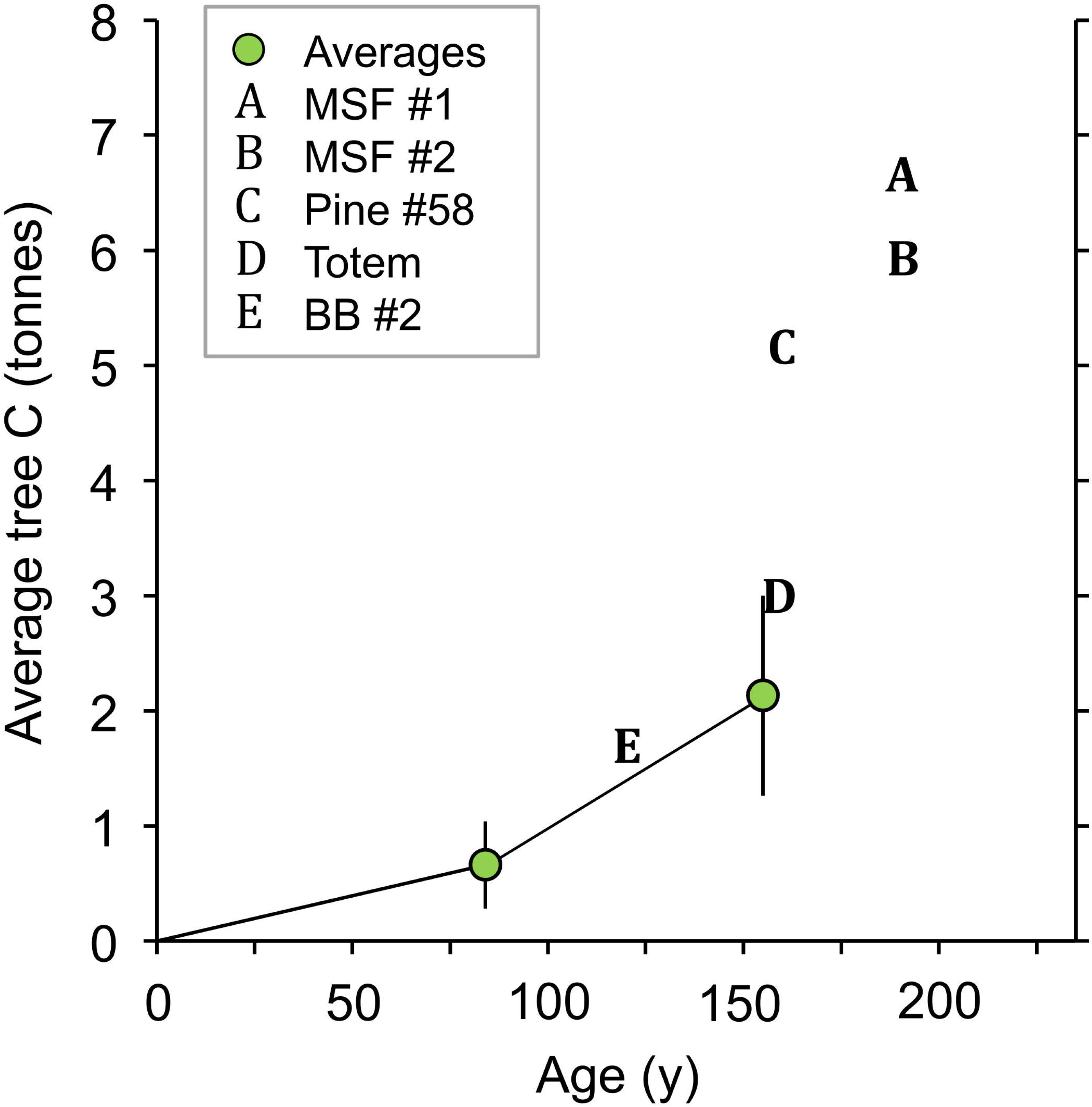
Tonnes of above-ground carbon (tC) in an “average” eastern white pine in a measured research acre (green locants) and in five individual trees (A,B,C,D,E) measured directly on site at three separate locations in Massachusetts. Average tC and standard deviation is based on pines in a stand at ~80 years (0.66 ± 0.38 tC) and ~160 years (1.95 ± 0.73 tC) as described in the text. Direct measurement of tC is shown for individual trees in western Massachusetts at these ages and locations: A, B - ~190 years (MSF #1 and #2, Monroe State Forest); C - ~160 years (Pine #58, Mohawk Trail State Forest; more details of Pine #58 shown in Figure 1); D - ~150 years (Totem, Northampton, MA); E – ~120 years (BB #2, Broad Brook, Florence, MA).

### 3.2 Stand measurements at ~80 and ~160 years

Detailed measurements were taken in comparable pine stands at ~80 and ~160 years *(TOP)*. As noted, the average tree in each stand is shown on Figure 2, and the distribution of tree sizes in the *TOP* is shown in Figure 3A. The largest pine in the *TOP* holds 4.24 tC and the smallest holds 0.53, an eight-fold difference. A comparison of the stand density and above ground carbon at ~80 vs. ~160 yr are shown in Figure 3B.

**Figure 3.**
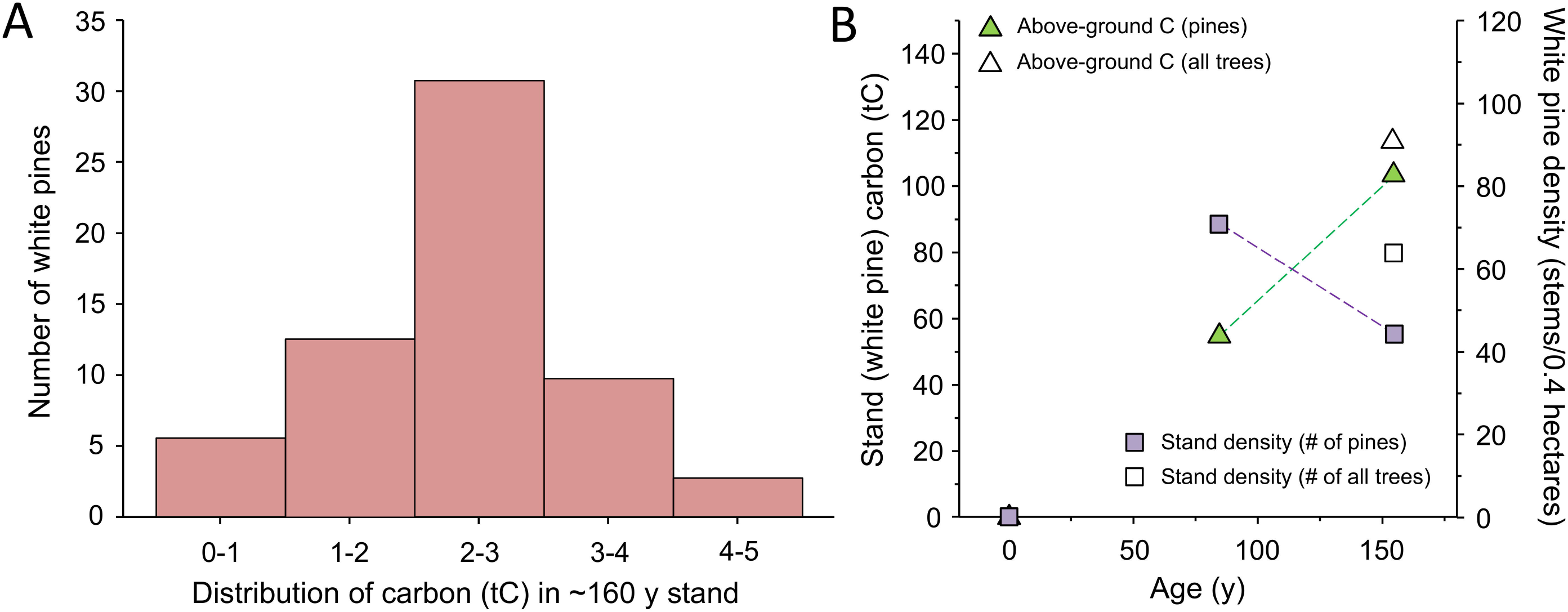
Carbon distribution, stand density and cumulative carbon in predominantly eastern white pine stands at ~80 and 160 years. These two stands were regrown from land previously used as pasture (i.e. not recovering from a harvest at time zero). **A.** Distribution of above-ground carbon (tC) among 76 eastern white pines of different sizes in the full *TOP* stand at ~160 years old. The majority contained 1-3 tC. **B.** Stand density and above-ground carbon measured directly on site in a research acre of eastern white pine at ~80 and 160 years. Stand density (# of stems) declined while above-ground carbon increased. The older stand includes some non-species that added to the number of stem and total carbon (open locants).

Complete data for 76 individual pines in the *TOP* (the 0.4 ha primary plot plus additional trees in the stand) is provided in Supplement 9. For comparison, data from 0.4 ha was collected from an ~80-year old stand growing on a terrace just downslope from the *TOP* in an area fairly well protected from wind and with adequate soil depth (Supplement 6). This age is more representative of the average stand of eastern white pine in New England. Average values for both ages are summarized in Table 1. As shown in Figure 2, we found an average of 0.66 tC per tree compared to 1.95 tC per tree in the *TOP*, a near tripling of carbon in the average individual pine in the older stand. We found a lower stand density in terms of number of stems, and a higher level of carbon in the *TOP*. Pines predominated both plots, and non-pine species added ~10% to the total above ground carbon (Figure 3B).

**Table 1.**
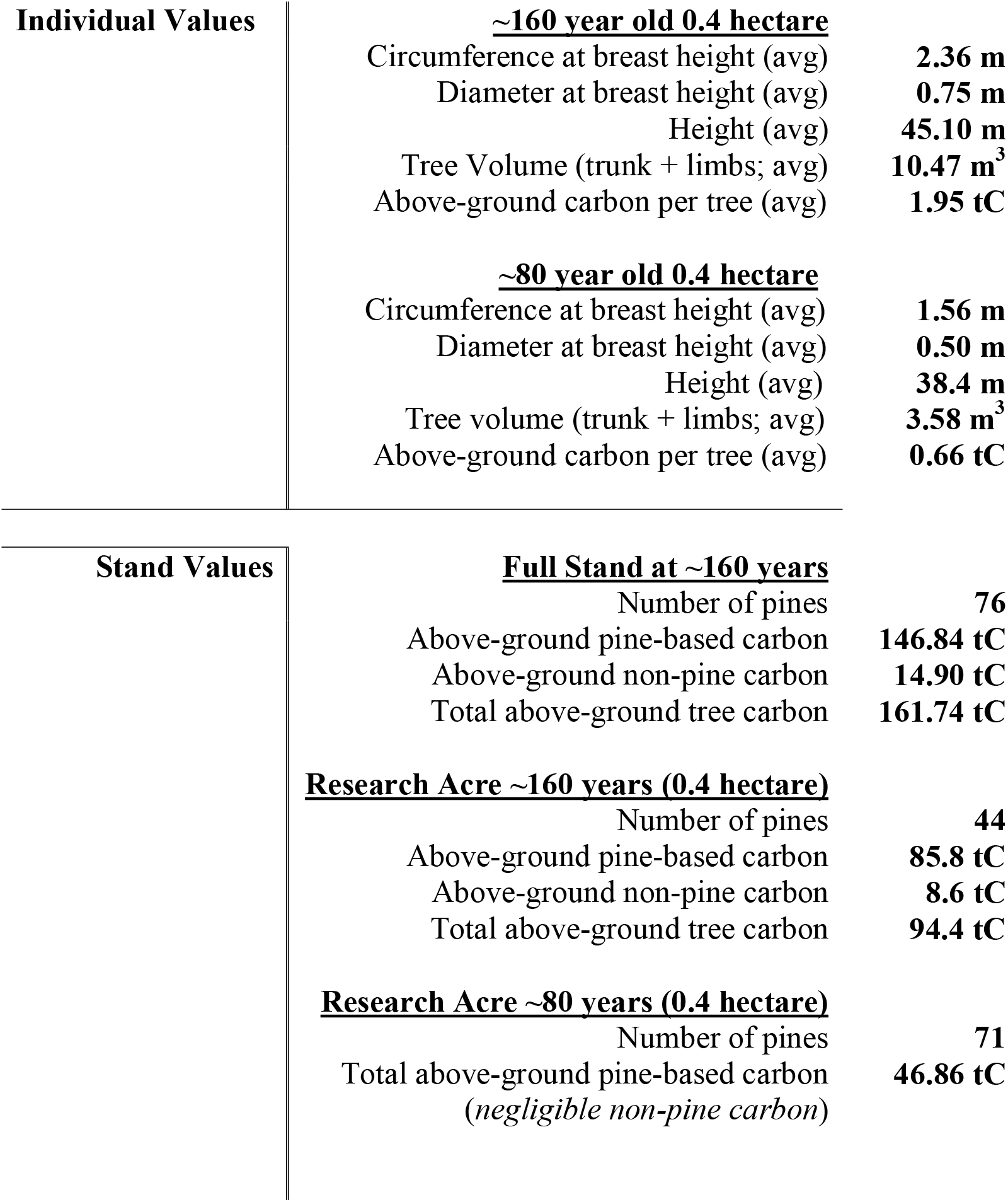
Summary of key measurements within a 160-year pine stand (*TOP*) and a comparable ~80 year old stand (2018 – 2019 values)

We note these calculations only include above ground tree-based carbon – they do not include more labile sources of additional carbon in the needles, leaves and understory plants, or the accumulation of woody debris in older stands. Our measurements also do not include the large store of underground carbon (the root system is typically estimated as an additional 15-20% of the above-ground tree volume, and total soil organic carbon can be an additional 50% or more. Therefore, the total carbon is significantly higher; we do not address those elements here. The above-ground tree-based carbon measured directly in the primary acre in the 80 year old stand is 46.86 tC and the 160-year-old stand is 94.4 tC, translating to 117.15 and 236.0 tC per hectare, respectively. Approximately 10% of the tree-based carbon in the older stand is non-pine; non-pine carbon in the younger stand is negligible (Table 1).

## 4. Discussion

The representative stands in this analysis approximate the average pine forest age (~80 years old) and a comparable stand nearly twice that age. To determine the biomass and above ground carbon in living trees as a function of tree size and age, we have used a combination of direct measurements and a hybrid FIA-COLE volume and biomass model to quantify individual trees and stands of eastern white pine, a common tree species in New England, We found that dominant individual trees accelerate their accumulation of carbon well past 150 years, and more than 75% of the carbon in dominant pines up to 190 years is gained after the first 50 years. Despite a lower stand density (fewer stems), total above-ground carbon is greatest in older stands and continues to increase past 150 years. The carbon per hectare quantified in these stands matches previous averages for the region and previous regional estimates that New England forests can accumulate at least twice as much carbon. The total carbon stored is even greater when below-ground carbon in roots, coarse woody debris, standing dead trees and smaller plants and soils are included (Nunery and Keeton, 2010; Tomasso and Leighton, 2014).

Forest managers stress the high accumulation rates of younger forests as important in absorbing atmospheric CO_2_. This is an important consideration for production forests to help optimize between growing a wood resource and accumulating carbon. Younger individual trees do not sequester more carbon than mature trees, and we did not find evidence for a significant benefit for a young stand compared to an older stand but we did not estimate accumulation rates below 80 years. Clearing an older forest to create a young forest creates a large carbon debt. Creating or maintaining this habitat has other benefits (wood production, habitat for hunting, or specific successional species), but it dramatically reduces forest carbon and eliminates the ability for that forest to host the full biodiversity of some of our rarest species of plants, animals, insects, fungi, lichens, reptiles and amphibians found in older and continuously forested areas (McMullin and Wiersma, 2019; Moose et al., 2019) as well as climate-sensitive birds that may benefit from old-growth forests (Betts et al., 2017). These older unmanaged forests also have fewer invasive species (Riitters et al., 2018).

The pine stands studied here grew from former sheep pastures, therefore not likely starting from a severely disturbed condition and thus minimizing initial carbon elution. This raises the question about the importance of site history in influencing growth, especially in the early years, since a disturbed condition can continue to lose carbon for more than a decade. We recognize that at some point above-ground carbon in living trees will no longer be increasing since the trees eventually die. However total forest carbon continues to increase even in some primary (“old-growth”) forests (Mackey et al., 2015). After tree death or forest disturbance there is a transfer of live carbon to dead wood and woody debris, the litter layer, and into the soil. Here in the older pine stand there is also the increased prevalence and growth of trees of other species (including more carbon-dense hardwoods) so that the species diversity and total carbon load continues to rise. A challenge for future research is to understand tree and stand carbon accumulation and dynamics in detail well beyond 200 years.

Public forests in New England are typically older than private forests (but still predominantly less than 100 years old), and provide the greatest possibility for intact forests across the landscape. Native tree species can live for several hundred years (and in the case of eastern hemlock (*Tsuga canadensis*) and black gum (*Nyssa sylvatica*), up to and exceeding 500 years). Despite the noted lack of old and old-growth forests, and the increasing level of natural disturbances from insects and storms creating forest diversity and forest openings, a major focus on public land is clearcutting to “create young forest” or “create resilience.” These programs assert that young forests prevent a suite of species from declining, that they sequester carbon more rapidly, and that they are more resilient than their older counterparts (Anwar, 2001). This approach has experimental merit, but at this time it needs more direct and long-term measurements and sufficient baselines and controls. It overlooks the dynamic evolution of these habitats over time, creating niches for these species. It also overlooks the critical role of cumulative stored carbon compared to sequestration, and the superior resilience of older forests to the stresses of climate change (Thom et al., 2019). The details of age and location (tropical, temperate, boreal, etc.) are also important in terms of what is reported “young” – in some cases considered up to 140 years (Pugh et al., 2019).

Our findings are consistent with Stephenson et al (2014) who found that absolute growth increases with tree size for most of 403 tropical and temperate tree species, and a study of 48 forest plots found that in older forests, regardless of geographical location, half of all above-ground biomass (and hence carbon), is stored in the largest 1% of trees as measured by diameter at breast height (Lutz et al., 2018). Keeton et al found an increase in carbon density per hectare as the age of the stand increased in the Northeast U.S. (Keeton et al., 2011) and a recent study in China found that forests with older trees and greater species richness had twice the levels of carbon storage than did less diverse forests with younger trees (Liu et al., 2018). Earlier work demonstrated that intact old growth forests in the Pacific Northwest contained more than twice the amount of sequestered carbon as did those that were harvested on a fixed rotation basis (Harmon et al., 1990). Erb et al. concluded that forests are capable of sequestering twice as much atmospheric carbon as they currently do (Erb et al., 2018). The potential for natural reforestation as a climate solution has just been increased dramatically (Cook-Patton et al., 2020).

Proforestation - growing existing natural forests - and recognizing the role of older forests and large trees in carbon accumulation and biodiversity protection, are critical components of a global strategy. Rapidly moving large stocks of atmospheric carbon as CO_2_ into forests and reducing emissions is essential for limiting the increase in global temperatures, and protecting intact and connected habitat is essential in preventing extinction. An important implication of this finding is that the estimated additional CDR achieved by future growth of secondary forests reported by Houghton and Nassikas is likely an underestimate because it does not account for ongoing accumulation rates as trees age (Houghton and Nassikas, 2018) – at least in regions with relatively young forests like those of the Northeast United States. The global study of natural forest carbon accumulation by Cook-Patton et al. provides quantitative evidence of the power of natural reforestation (Cook-Patton et al., 2020). Considering these reports and the current findings increases the potential regional contribution for increased carbon accumulation rates in the coming decades by Northeastern temperate forests. While the IPCC clearly identified forests as essential for sequestering additional carbon for climate stability it focused on production forests that are currently recovering from being harvested or on unforested areas where forests could be planted (afforestation). A report by Bastin et al. proposes massive afforestation on 0.9 billion ha but acknowledges that it will take time before large amounts of carbon would be sequestered (Bastin et al., 2019). Global tree planting efforts are under way but there is little data on how to plant an ecosystem, and some of these tree planting efforts suffer from 75-80% mortality of the young trees. In contrast, growing existing forests that already contain large carbon stores and can rapidly sequester increasing amounts of atmospheric carbon dioxide and accumulate diversity over time - a much more effective near-term and proven strategy (Moomaw et al., 2019). While valuable, neither afforestation nor reforestation will remove as much atmospheric carbon as proforestation in the next 50 years – the timeline when it is needed most to avoid irreversible consequences of a changed climate. Protecting primary forests and secondary forests where possible is also a far better option than the unproven technology of bioenergy with carbon capture and storage (BECCS), also suggested by the IPCC report (Anderson and Peters, 2016; IPCC, 2018). Finally, letting existing secondary forests grow provides equity, natural heritage and cumulative health benefits for people in terms of respite and passive recreation - and does not compete directly with agriculture and other demands for land use.

The highly accurate direct measurements at the tree and stand level in this paper are consistent with parameterized and other studies at larger scale in verifying that larger trees (Lutz et al, 2018, Stephenson et al, 2014) and stands of larger trees accumulate the most carbon over time compared to smaller trees (Mildrexler et al. 2020). They support the proforestation strategy of growing existing forests to achieve their natural capacity to accumulate carbon and achieve their biodiversity potential (Moomaw et al., 2019) to redress the balance of carbon lost to the atmosphere from global forests (Hudiburg et al., 2020). The important implication of these findings is that the trees and the forests that we need most for carbon storage and CDR to help limit near-term climate change are the ones that are already established.

Plantations and forests managed for forest products account for 71% of all forest area globally (IPCC, 2019), more than sufficient for resource production. Strategic planning can enable some to be prioritized and repurposed for climate protection and research - and the remaining 29% should be protected wherever possible. High levels of carbon accumulation and biodiversity protection can be achieved in parallel with inherent resiliency to a changing climate – including by protecting species networks, genetic diversity and epigenetic changes. These findings also specifically ground-truth the capacity for New England pine forests to more than double their carbon in the coming decades. Protection of public forests from unneeded intervention is urgent, and compensation programs should be established for stewarding private forests based on numerous ecosystem services.

## Acknowledgments

We acknowledge the critical technical assistance of Jared D. Lockwood in measuring pines in the *TOP*, the younger pine stand, and elsewhere as needed. We thank Monica Jakuc Leverett for invaluable assistance in editing and revising the original draft, and David Ruskin for tireless efforts throughout the process. We thank Ray Asselin for photographing pines for further analysis, and for ring and whorl counts to establish ages. Supported by Trinity College and a Charles Bullard Fellowship in Forest Research (SAM), a Faculty Research Grant from the NASA Connecticut Space Grant Consortium (RTL, SAM) and the Rockefeller Brothers Fund (WRM).

## Author contributions statement

RTL chose site locations and individual trees, established measurement methods and protocols, did the on-site tree measuring, and performed the subsequent analysis. SAM analyzed and organized the content and supplements and participated in drafting and finalizing the text. WRM framed the analysis in the context of other studies and the larger context of climate change, assisted with data analysis and presentation, and drafting and editing the text.

