## Supplements for "Older eastern white pine trees and stands sequester carbon for many decades and maximize cumulative carbon"

**Supplement 1**

**Native Tree Society exceptional and/or well-documented white pine sites**

The Native Tree Society (NTS) has studied many white pines sites over 25 years, confirming the species maximum size across a wide variety of site conditions and age ranges.

1. Mohawk Trail State Forest, MA
2. Monroe State Forest, MA
3. Ice Glen, Stockbridge, MA
4. Harvard Forest, Petersham, MA
5. William Cullen Bryant Woods, Cummington, MA
6. Upper Broad Brook, Northampton, MA
7. Look Park, Northampton, MA
8. Carlisle Pines, Carlisle, MA
9. Cathedral Pines, Cornwall, CT
10. Cook Forest State Park, PA
11. Hearts Content Natural Area, PA
12. Anders Run State Park, PA
13. Delaware Water Gap, PA
14. Elders Grove, Adirondack Park, NY
15. Pack Forest, Adirondack Park, NY
16. High Peaks Wilderness Area, Adirondack Park, NY
17. Five Ponds Wilderness, Adirondack Park, NY
18. Lily Dale, NY
19. Ordway Pines, Norway, ME
20. Bowdoin College Pines, ME
21. Hemenway State Forest, NH
22. Bradford Pines, NH
23. Private site, Claremont, NH
24. Pine Park, Dartmouth College, NH
25. College Pines, Hanover, NH
26. Marsh-Billings-Rockefeller National Historical Park, Woodstock, VT
27. Fisher-Scott Memorial Pines, Arlington, VT
28. Great Smoky Mountains National Park, TN-NC (multiple locations)
29. Linville Gorge Wilderness, NC
30. Cullasaja Gorge, NC
31. Ellicott Rock Wilderness, GA/NC/SC
32. Cooper’s Creek Wildlife Management Area, GA
33. Cohutta Wilderness, GA
34. Ramsey’s Draft Wilderness Area, VA
35. Porcupine Wilderness State Park, MI
36. Sylvania Wilderness, MI
37. Hartwick Pines State Park, MI
38. Lake Itasca State Park, MN

**Supplement 2**

**NTS direct measurement methodology used for determining volume**

**Introduction**

A critical method used in calculating the volume of white pine trunks in this paper employs a guide by Native Tree Society (NTS). We call the method direct measuring to distinguish it from calculating volumes through allometric equations, which are statistical derivations. A detailed description is presented in Leverett et al. (2020); a summary is offered here.

**The process**

Measuring a tree’s trunk for volume involves dividing it into adjacent sections and measuring the diameters of the top and base and the height of each, typically employing the combination of tape, laser hypsometer, tripod, and monocular with range-finding reticle. The values obtained for a section are entered into a formula for a geometric frustum, commonly a neiloid, paraboloid, or cone. The frustums usually have circular cross-sections, but can be elliptical. The volumes of the individual sections are added to get the trunk volume. A number of Excel workbooks that automate the calculations are available on the NTS website (www.nativetreesociety.org) and at FEMC (www.uvm.edu/femc). Formulas used in the modeling process are summarized below.

**Basic frustum volume formulas**

Variable definitions

V = frustum volume

H = frustum height

r_1_ = radius of base of frustum

r_2_ = radius of top of frustum

1. Right circular cone: $V= \left( \frac{1}{3} \right)\pi H\left( r_{1}^{2}+r_{1}r_{2}+r_{2}^{2} \right)$
2. Right circular paraboloid: $V= \left( \frac{1}{2} \right)\pi H\left( r_{1}^{2}+r_{2}^{2} \right)$
3. Right circular neiloid: $V= \left( \frac{1}{4} \right)\pi H\left( r_{1}^{2}+r_{1}^{\frac{4}{3}}r_{2}^{\frac{2}{3}}+r_{1}^{\frac{2}{3}}r_{2}^{\frac{4}{3}}+r_{2}^{2} \right)$

The Excel workbook that automates this modeling process allows the user to select the type of frustum to use for each section. Advanced versions allow frustums of intermediate forms to be used as well as frustums with elliptical cross-sections. Other refinements provide compensations for head swivel when a typical photographer’s tripod is used to steady the laser hypsometer.

**Reticle formula**

The monocular with range-finding reticle and the hypsometer are the instruments used to measure trunk diameter from a distance. While there are several formulas that can be used, the primary one is shown below.

Variable definitions

W = trunk width (diameter)

M = reticle reading

D = distance to front middle of trunk

F = manufacturer’s reticle factor

$$W= \frac{MD}{F-0.5M}$$

**Height formula**

The laser hypsometer is used to measure heights on a trunk. There are usually two main height routines that implement the sine and tangent methods. The primary routine we use is the sine method that implements the following formula:

Variable definitions

H = height being measured

L_1_ = hypotenuse distance to lower point on trunk

L_2_ = hypotenuse distance to upper point on trunk

A_1_ = angle to lower point on trunk

A_2_ = angle to upper point on trunk

$$H=L_{2}sin\left( A_{2} \right)-L_{1}sin\left( A_{1} \right)$$

In this formula, the sine of a negative angle is negative.

**Summary**

The above explanation outlines only the basic process. Where trunk visibility is a problem, or the trunk shape differs significantly from circular, refinements to the basic process are employed, which may require photographs of the trunk to aid in identifying the base and top of each frustum as seen from different sides of the trunk. Measurements from perspectives 90° apart are useful to determine circularity or lack thereof.

**Supplement 3**

**Use of form factor in measuring trunk volume of individual trees**

**Introduction**

The USFS uses statistically derived trunk form and taper factors in their allometric equations to compute trunk volumes. The methodology is well established. These factors allow bole volumes to be calculated from tree height and diameter inputs, or just diameter. The factors are broad statistical averages. In this paper we employ one of the Forest Service Inventory and Analysis (FIA) models, and we also derived trunk form factors specifically applicable to the mature white pines in the *Trees of Peace* (*TOP*) based on volume-modeling (see Supplement 2). We also explored trunk form factor values that could be assigned white pines across the range of sizes in the *TOP*, appealing to traditional trunk taper models. These three tools allow us to more confidently calculate trunk volumes in the field using diameter at breast height (DBH) and full tree height. A detailed description is presented in Leverett et al. (2020); a summary is offered here.

**Composite form factor based on neiloid, cone, and paraboloid averaging**

Traditionally, the trunk of a conifer such as a stand-grown white pine has been visualized as encompassing three rates of taper. From the base up to between one and two m, the trunk follows a concave shape, which can be described as a neiloid. Above that, the trunk taper is slightly convex, suggesting the shape of a paraboloid, and finally a straight conical taper. In terms of the volumes treated as three-dimensional solids, we have the general formula:

$$V=FH\pi\frac{D^{2}}{4}$$

where F = trunk form factor

H = full trunk height

D = diameter at breast height

This formula can also be thought of as giving the part of a cylinder occupied by the trunk, where the height of the cylinder is H and the base of the cylinder has an area $\pi\frac{D^{2}}{4}$. The value of F specifies the portion of the cylinder occupied by the trunk. For a concave neiloid-shaped trunk,

F = 0.25, 0.5 for a convex paraboloid, and 0.333 for a straight-sided cone. Can we arrive at an average value for F that would apply to the whole trunk? On white pines in the *TOP*, from the base up to 1.37 m can be treated as neiloid. The start of the crown is usually between 27 and 30 m for trees in the height range of the average height of a pine of 45.1 m. We will use 28.5 m. We assign the percentages of height for the three factors of

N = 0.03

P = (28.5 - 1.37)/45.1 = 0.602

C = (45.1-28.5)/45.1 = 0.368

The weighted factor is then 0.03 x 0.25 + 0.602 x 0.5 + 0.368 x 0.333 = 0.4310, a result surprisingly close to 0.4303, obtained by the averaging of the form factors calculated in

Table S3.1. This result supports the continued relevance of the simple neiloid-paraboloid-cone trunk model.

**Average form factors from direct measurement and FIA-COLE model**

A sample of 39 white pines ranging in trunk volume from 30.81 m^3^ down to 1.69 m^3^ were determined through the NTS direct measuring approach (Supplement 2) and by the FIA-COLE model (Supplement 4). The results are shown below. For more detailed data, see Table S3.2.

**Table S3.1 Comparison of trunk form factors**

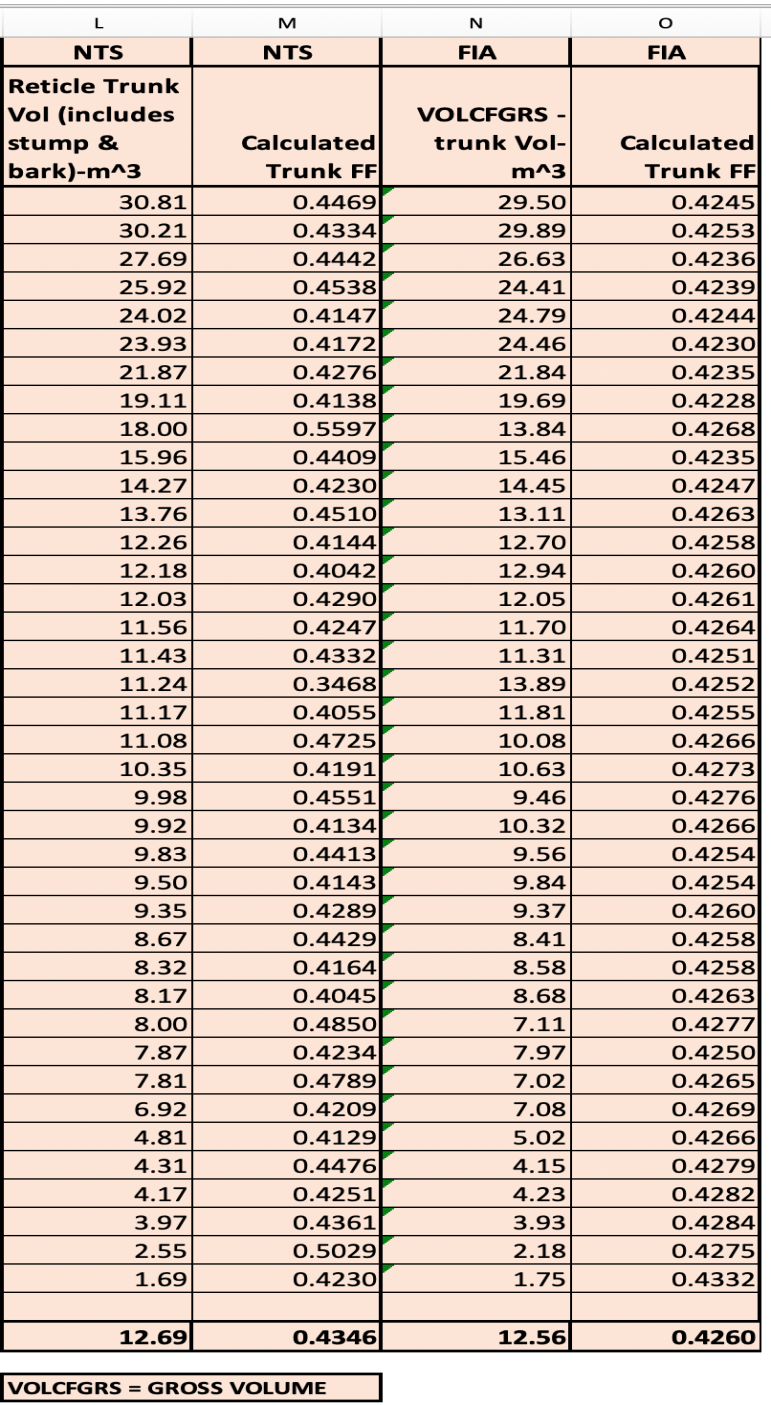

The FIA-COLE model does not use form factors directly. However, once a volume is determined, its effective form factor can be computed for comparison to the NTS counterpart

using	$F= \frac{4V}{\pi HD^{2}}$

The FIA-COLE model’s average form factor is 0.4260, the NTS model yields 0.4346, and the average of the two is 0.4303. The principal difference in the two methods is that direct measurement is sensitive to deviations from the norm, and the FIA-COLE model reflects that statistical norm (Leverett et al., 2020)

The NTS form factor differs from FIA-COLE by less than 2.0%.

Final carbon values were derived by averaging the NTS and FIA-COLE results. We chose the averaging process to recognize the value of both methods of computing carbon in the trunks of live pines.

Our analysis led to one other surprising result. In the FIA-COLE model, the trunk volume can be closely approximated with the equation:

$\boldsymbol{V=0.33363}\boldsymbol{H}\boldsymbol{D}^{\boldsymbol{2}}$

where H = full height of the tree and D = diameter at breast height. The model and its approximation differ by an average of less than 0.1%. This simple formula offers foresters and forest scientists an extremely easy way to calculate trunk volume when in the field.

A second formula can be derived that adds in crown volume for white pines in the age range of the tree being studied. From Supplement 5, if we use a 15% limb factor, we get a full trunk and limb volume (tree volume, or total above-ground volume) formula of:

$$\boldsymbol{V=0.38367}\boldsymbol{H}\boldsymbol{D}^{\boldsymbol{2}}$$

These two formulas are applicable to stand-grown white pines.

**Table S3.2**

**39 pines – detailed reticle measurements compared to FIA-COLE model**

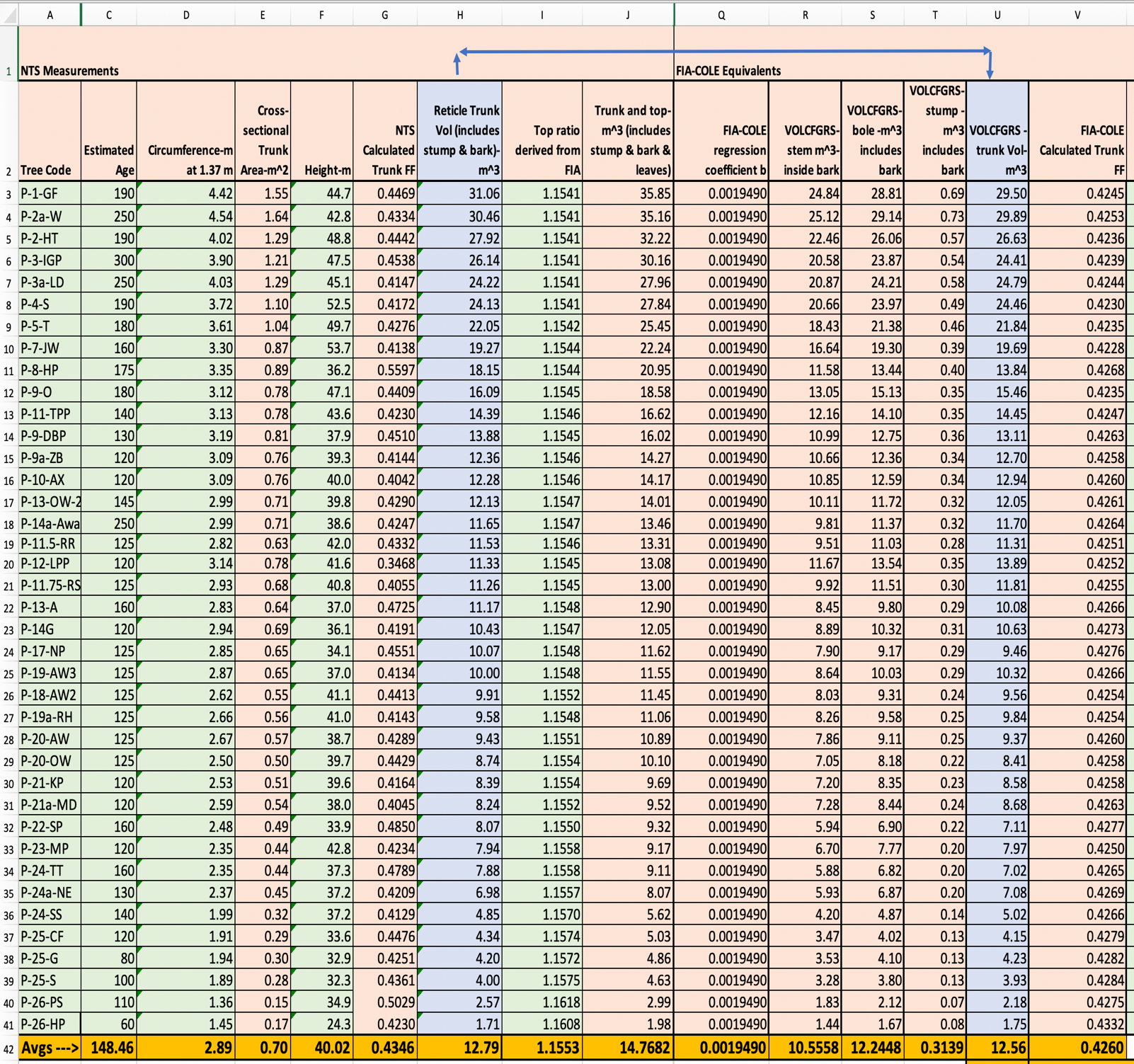

**Supplement 4**

**FIA-COLE volume-biomass model for trunk volume**

We chose an FIA-COLE based volume and biomass model offered as a COLE tool to compute biomass in individual trees of 189 species of which white pine is one. We employed the model as a parallel and alternative method to direct measurements for computing trunk and limb volume and their associated mass equivalents. We then took the average of the two models. Here we define variables used in the model, which uses Imperial units.

1. **Variables**

V_s_ = volume of stem (wood only, no bark) from a 1-foot stump to a 4-inch top measured to outside bark. This is defined in the FIA database as VOLCFGRS, which stands for gross cubic foot volume of the stem.

b = regression-based value for white pine = 0.001949

V_b_ = volume of bole [volume of stem + volume of bark (bark is 0.16 x V_s_)]

V_st_ = volume of stump, outside bark (this is bark and wood)

V_t_ = volume of trunk to a 4-inch top (trunk = bole + stump)

M_st_ = mass of stump in lbs

D = diameter at 4.5 feet above mean base level in inches (41.37 inches, Pine #58)

H = full height (176.2 feet, Pine #58)

2. **Equations**

D is in inches and H is in feet.

A. V_s_ = b(D^2^H) = 0.001949D^2^H. (regression-based)V_b_ = 1.16V_s_ (Jenkins bark factor)
B. V_st_ =0.005454153D^2^ [1+(5.62462)(0.08091)+8.50038 (0.08091^2^)]. (Raile stump volume)
C. V_t_ = V_b_ + V_st_ (trunk volume to a 4-inch top – no limbs)
3. **Calculations for Pine #58**:

V_s_ = 0.001949[41.37^2^(176.2)] = 587.74 ft^3^

V_b_ = 1.16V_s_ = (1.16)(587.74) = 681.78 ft^3^

V_st_ = 0.005454153(41.37)^2^ [1+(5.62462)(0.08091)+8.50038 (0.08091^2^)] = 14.10 ft^3^

V_t_ = 681.78 + 14.10 = 695.88 ft^3^

**Supplement 5**

**Derivation of FIA-COLE limb factor**

The limb-to-trunk ratios that we use are a derived result from the FIA Jenkins and Raile models using biomass as a surrogate for volume. The model computes total biomass, stem wood, stem bark, foliage, root, stump, and top biomass. By definition, the top includes the limbs, branches, and twigs, but not foliage. This is what we want to compare to our direct measurements of above-ground carbon. For large trees, the range is generally between 15% and 16%, with the percentage going up as DBH goes down. The model is sensitive only to DBH and reaches a maximum at 5 inches DBH of 21.4%. We employed the model down to a diameter of 7.92 inches, which gave an 18.5% limb factor.

The FIA-Jenkins-Raile model’s top and foliage calculations are as follows:

(AG = Above Ground; B1 and B2 are factors used by the model)

Total_AG_biomass_Jenkins (lbs) = Exp(Jenkins_Total_B1 + Jenkins_Total_B2 * ln( DBH × 2.54 )) × 2.2046

Jenkins_Total_B1 = -2.5356

Jenkins_Total_B2 = 2.4349

Total_AG_biomass_Jenkins (lbs) = Exp(-2.5356+ 2.4349* ln( DBH × 2.54 )) × 2.2046

Foliage Ratio = EXP(-2.9584+(4.4766/(2.54DBH))) where DBH is in inches

Jenkins Foliage biomass = Jenkins Foliage Ratio x Total AG biomass Jenkins

Top (limbs) = total above ground biomass – bole biomass – foliage biomass – stump biomass

Trunk biomass = bole biomass + stump biomass (our definition)

Using Pine #58, featured in Figure 1, we have the following (biomass figures taken from Excel workbook implementing the model):

Total AG biomass Jenkins (lbs) = 14,599.23

Foliage Ratio = EXP(-2.9584+(4.4766/(2.54(41.37))))= 0.0543

Jenkins Foliage biomass 0.0543 x 14,599.23 = 792.73

Top (limbs) = 14,599.23 – 11,647.76 – 792.73 – 313.97 = 1,844.77 lbs

Biomass ratio limbs to trunk = 1,844.77/(11,647.76 + 313.97) = 0.1542

We used this biomass ratio to substitute for the volume equivalent.

**Supplement 6**

**Analysis of above-ground carbon in an ~80 y-old stand**

An important part of the process was to study a stand of young pines to develop an independent profile for comparison to the *TOP*. We identified a site in Mohawk Trail State Forest a short distance from the older trees. The following steps summarize our approach:

1. Within the selected site, we chose a sample of pines and counted annual whorls up to visible heights to make age estimates. We included the largest and smallest pines in the stand to establish the size range. We measured the heights and DBHs of the sample pines.
2. We established a densely populated point-centered subplot with a 12.2 m radius that included 12 pines for concentrated measuring. The area of the subplot equaled 467.7 m^2^. Projecting density to a 0.4-ha area, we have 12(4046.9/467.7) = 104 pines.
3. We counted the number of white pines in two separate 0.4-ha plots in the stand (a standard acre). The count was 70 stems for the first plot and 71 for the second. Beyond the boundaries of the plots, the stand quickly thins out. So, we set average density at **71 stems** in the 0.4-ha plot with a high of 104 for a dense plot.

4. The largest pine has a 2.08-m circumference. For this pine, we counted 40 whorls (equals 40 years) to a height of 23.16 m. The height was 40.6 m in height, with an average 0.58 m per year over the 40 y. The remainder of the tree, 17.4 m, was assigned a growth rate of 0.4 m/y - probably high, but this tree was one of the tallest. This resulted in a projected age of 84.0 y.
5. For a second 39.6 m tall pine, we counted 44 whorls up to a height of 24.1 m or 0.55 m/y. The projected age was:

44 + (39.6 - 24.1)/0.4 = 82.8 y

6. Other computed internode lengths calculated by the ratio of height/# whorls gave:

12.53/25 = 0.5 m

10.67/21 = 0.51 m

13.72/25 = 0.55 m

19.57/34 = 0.58 m

15.42/25 = 0.62 m

15.79/25 = 0.63 m

The average is 0.564 m, which almost exactly matches the average of the first two trees, which were 0.58 and 0.55 m.

7. As a check on the lower trunk internode growth, we measured the leader (highest internode on the trunk) of a conspicuous pine to 0.28 m. Other pines showed annual growth in the range of 0.24 to 0.40 m, suggesting that our 0.40-m projection of internode length beyond the bole heights of the whorl counts to be high. If so, our stand age of 84 y is low. However, the development of the stand likely occurred over a 10 to 15-y period and we were not able to collect any data to suggest an older stand age (i.e. beyond 84 y).
8. We measured heights for all 12 pines in the subplot to the highest level of accuracy we could attain, using the LTI TruPulse 200X’s built-in sine method. We got the following range of heights:

35.2, 35.8, 36.1, 36.2, 37.1, 37.1, 39.6, 40.5, 40.6, 40.6, 40.9, 41.1

The average is **38.4 m** and range is 5.9 m (standard deviation 2.3), which is about half the range we documented within the *TOP*. Younger stands are generally more uniform in height.

9. The average circumference of the 12 pines was 1.56 m with a minimum of 0.77 and maximum of 2.09 m. We chose an average trunk form factor **of 0.4346** (derived from complete volume modeling of 39 pines; Supplement 1), reflecting the age of the pines.

With the 1.56, 38.4, and 0.4346 factors, we used equation S2.1 as follows

$$V=\frac{C^{2}}{4\pi}HF$$

where C = circumference at 1.37 m

H = total tree height

F = trunk form factor

Applying this formula gives

$V=\frac{{1.53}^{2}}{4\pi}\left( 38.4 \right)0.4346=3.11$ m^3^

Applying the 1.15 limb multiplication factor (Supplement 5), we arrived at an average 3.58 m^3^. In terms of carbon, this translates to 0.66 tC per tree.

As a check on maximum annual growth, the biggest pine in the cluster of 12 pines used as a subsample measured 2.08 m in girth. Its radius equals 0.33 m. The average ring width is 0.39 cm. With a height of 40.6 m, the estimated volume of this pine is

V= π(0.33)^2^(40.6)(0.4346) = 6.04 m^3^

Adding 15% for limbs, gives 6.95 m^3^, which translates to 1.29 tC.

**Supplement 7**

**Pine #58’s measurement history**

The *TOP* include Pine #58. As of 2020, Pine #58 is the tallest accurately measured tree in New England, and possibly the Northeast. It is also the largest pine by volume in the *TOP*.

Pine #58 was initially measured with a transit by Jack Sobon and Bob Leverett in November 1992. The height at that time was 47.24 m, and the DBH was 0.94 m. The pine was climbed in 1998, 2001, and 2004 to determine its tape-drop height and to volume-model its trunk.

Pine #58 has been measured for height from the ground by at least a half dozen NTS members using a variety of instruments. Its height of 53.71 m and diameter of 1.05 m is well established.

To arrive at the current measurement of trunk volume for Pine #58 using the NTS method, we used its height of 53.71 m, DBH of 1.05 m, and a trunk form factor of 0.4346 (Supplement 2). This gave us a total trunk volume of 24.25 m^3^.

The total wood and bark volume of the above ground part of Pine #58 includes limbs, branches, and twigs. The FIA-COLE volume and biomass model gives a limb factor as applied specifically to Pine #58 is 1.1544. This gives us 27.99 m^3^ and the equivalent carbon is 5.18 tC.

Although we settled on the average 1.15 limb volume factor, we tested its applicability to Pine #58 using a separate limb profile. We made the following assumptions:

1. each limb whorl has 4 branches
2. maximum crown radius for all whorls = 7.92 m
3. radius at right angles to the maximum = 6.73 m (the two produced a maximum average crown spread of 14.65 m, matching what we measured from the ground)
4. maximum limb radius at top of crown = 0.61 m
5. radius at right angles to the top = 0.52 m
6. depth of the crown (23.16 m)
7. maximum limb diameter (11.43 cm)
8. form factor for limb of 0.4167 (midway between cone and paraboloid)
9. 10% of limb volume extra to cover branches and twigs

Using an arithmetic progression from base to top, the volume of limbs and branches came to

2.81 m^3^. Using an NTS-FIA average trunk volume of 19.97 m^3^, we get a limb-to-trunk ratio of (2.81/19.97) = 0.14. The FIA-COLE model gives a range of limb factors for the pines we included in the study, averaging 0.16 for the *TOP*. We averaged the 0.14 and 0.16 and settled on 0.15 as an acceptable factor for the *TOP* as a whole.

**Supplement 8**

**Young versus old pines - equivalent radial growth**

Young pines often exhibit visibly prominent growth in the width of their rings and in the length of their annual internodes. For example, a young pine exhibiting 0.61 to 0.91-m internode lengths on the trunk suggests that the significant growth comes in the early years of life – which it does in terms of these two dimensions. However, once a pine has gained considerable trunk volume, a small added ring and modest new internode length can represent a much larger volume gain than is visually apparent.

On simple geometric models, such as a cone, doubling both the height and diameter leads to an 8-times volume increase. If the form factor and limb factors change, the multiplier may go as high as 10. See Table S9, below.

**Table S9 Comparing volumes by doubling diameter and height**

|  | Diameter(m) | Height (m) | Form Factor | Volume (m^3^) | Limb Factor | Total Volume (m^3^) |
| --- | --- | --- | --- | --- | --- | --- |
| 1 | 0.3 | 24 | 0.4346 | 0.74 | 1.15 | 0.85 |
| 2 | 0.6 | 48 | 0.4346 | 5.90 | 1.15 | 6.79 |

This hypothetical tree doubles in diameter and height. The tree’s form and limb factors stay the same, but these changes cause the volume to increase by a factor of 8. The significance of this is that **a small trunk radial increase in a large tree can match the volume growth of a smaller tree growing more rapidly in terms of annual radial and height increase**.

We derived a useful formula to compute the increase in radius needed for one tree of known dimensions and height increase to match the volume change in a second one. The full dimensions of the second tree and its height and radial changes are also known. See equation below.

$$\Delta r_{1}=\sqrt{\frac{F_{1}r_{1}^{2}h_{1}+F_{2}\left( r_{2}+\Delta r_{2} \right)^{2}\left( h_{2}+\Delta h_{2} \right)-F_{2}r_{2}^{2}h_{2}}{F_{1}\left( h_{1}+\Delta h_{1} \right)}}-r_{1}$$

where	r_1_	= radius of large tree, and r_2_ = radius of small tree

h_1_	= height of large tree, and h_2_ = height of small tree

F_1_	= form factor for large tree, and F_2_ = form factor for small tree

∆r_1_ = change in radius for large tree, and ∆r_2_ = change in radius for small tree

∆h_1_ = change in height for large tree, and ∆h_2_ = change in height for small tree.

**Supplement 9**

**Measurements of the *Trees of Peace***

Here we report the heights, diameters, and trunk and limb volumes of the 76 pines in the primary stand of the *TOP.* 44 of these pines are in the measured 0.4 ha (1 acre) plot. Some pines have been measured multiple times over years including Pine #58, dating back to 1992.

27 pines in the *TOP* reach or exceed 45.72 m in height (approximately 150 ft) and another 15 reach this threshold in adjacent areas. In Mohawk Trail State Forest at least 146 pines meet or exceed 45.72 m.

The measurements in Table S9 below represent our volume determinations for the 76 white pines that are alive and growing in the *TOP*. The average carbon mass per pine is 1.93 tC.

**Table S9 Measurements of *TOP***

The following table lists measurements of the 76 pines in the *TOP*. DBH, height, and form factor (derived from the 39-pine sample - Supplement 1) were used to calculate NTS volume. The FIA-COLE model was applied using DBH and height.

| **Measurements of the TOP – summary** | | | | | |
| --- | --- | --- | --- | --- | --- |
| Tag # | DBH (m) | Standard Trunk Form Factor | Height (m) | Avg NTS & FIA Volume-trunks & limbs (m^3^ ) | Avg NTS & FIA Carbon -trunks & limbs (tonnes) |
| 58 | 1.05 | 0.4346 | 53.71 | 23.00 | 4.24 |
| 77 | 1.09 | 0.4346 | 47.58 | 21.83 | 4.03 |
| 27 | 1.08 | 0.4346 | 47.24 | 21.25 | 3.92 |
| 52 | 0.96 | 0.4346 | 51.51 | 18.37 | 3.39 |
| 26 | 0.95 | 0.4346 | 48.77 | 17.13 | 3.16 |
| 105 | 0.98 | 0.4346 | 44.81 | 16.74 | 3.09 |
| 104 | 0.98 | 0.4346 | 44.81 | 16.63 | 3.07 |
| 38 | 0.94 | 0.4346 | 48.04 | 16.46 | 3.04 |
| 5 | 0.95 | 0.4346 | 45.78 | 16.23 | 2.99 |
| 65 | 0.99 | 0.4346 | 41.00 | 15.54 | 2.87 |
| 63 | 0.92 | 0.4346 | 46.63 | 15.28 | 2.82 |
| 90N | 0.91 | 0.4346 | 44.81 | 14.44 | 2.66 |
| 111 | 0.91 | 0.4346 | 44.81 | 14.44 | 2.66 |
| 41 | 0.88 | 0.4346 | 46.79 | 13.93 | 2.57 |
| 55 | 0.89 | 0.4346 | 44.81 | 13.85 | 2.56 |
| 73 | 0.91 | 0.4346 | 42.64 | 13.85 | 2.56 |
| 25 | 0.89 | 0.4346 | 44.81 | 13.81 | 2.55 |
| Missing | 0.86 | 0.4346 | 45.87 | 13.35 | 2.46 |
| Missing | 0.85 | 0.4346 | 45.93 | 13.00 | 2.40 |
| 31 | 0.83 | 0.4346 | 47.88 | 12.88 | 2.38 |
| 34 | 0.83 | 0.4346 | 47.12 | 12.76 | 2.36 |
| 48 | 0.81 | 0.4346 | 49.35 | 12.72 | 2.35 |
| 68 | 0.88 | 0.4346 | 41.61 | 12.49 | 2.30 |
| 37 | 0.82 | 0.4346 | 47.27 | 12.39 | 2.29 |
| 35 | 0.82 | 0.4346 | 46.88 | 12.27 | 2.26 |
| 71 | 0.83 | 0.4346 | 44.81 | 11.96 | 2.21 |
| Missing | 0.83 | 0.4346 | 44.04 | 11.80 | 2.18 |
| 95 | 0.82 | 0.4346 | 43.53 | 11.52 | 2.13 |
| 49 | 0.79 | 0.4346 | 47.00 | 11.48 | 2.12 |
| 66 | 0.81 | 0.4346 | 44.81 | 11.38 | 2.10 |
| 36 | 0.80 | 0.4346 | 45.57 | 11.22 | 2.07 |
| 79 | 0.80 | 0.4346 | 44.93 | 11.22 | 2.07 |
| 50 | 0.80 | 0.4346 | 44.81 | 11.22 | 2.07 |
| 56 | 0.80 | 0.4346 | 44.81 | 11.22 | 2.07 |
| 50 | 0.79 | 0.4346 | 46.63 | 11.20 | 2.07 |
| 74 | 0.80 | 0.4346 | 44.20 | 11.07 | 2.04 |
| 70 | 0.78 | 0.4346 | 46.79 | 10.96 | 2.02 |
| 29 | 0.79 | 0.4346 | 44.81 | 10.95 | 2.02 |
| 51 | 0.81 | 0.4346 | 42.43 | 10.71 | 1.98 |
| 28 | 0.76 | 0.4346 | 44.81 | 10.09 | 1.86 |
| 44 | 0.73 | 0.4346 | 47.64 | 9.93 | 1.83 |
| 72 | 0.75 | 0.4346 | 44.81 | 9.86 | 1.82 |
| 85 | 0.74 | 0.4346 | 45.84 | 9.70 | 1.79 |
| 78 | 0.73 | 0.4346 | 44.81 | 9.33 | 1.72 |
| Missing | 0.72 | 0.4346 | 46.21 | 9.26 | 1.71 |
| 28 | 0.72 | 0.4346 | 45.84 | 9.19 | 1.70 |
| 39 | 0.72 | 0.4346 | 44.81 | 9.11 | 1.68 |
| 113 | 0.72 | 0.4346 | 44.81 | 9.11 | 1.68 |
| 80 | 0.72 | 0.4346 | 44.81 | 9.01 | 1.66 |
| 90 | 0.71 | 0.4346 | 44.81 | 8.86 | 1.64 |
| 62 | 0.71 | 0.4346 | 44.81 | 8.70 | 1.61 |
| 39 | 0.72 | 0.4346 | 42.06 | 8.44 | 1.56 |
| 40 | 0.69 | 0.4346 | 45.14 | 6.89 | 1.27 |
| 45 | 0.67 | 0.4346 | 46.63 | 8.25 | 1.52 |
| 76 | 0.68 | 0.4346 | 44.20 | 7.89 | 1.46 |
| 108 | 0.67 | 0.4346 | 44.20 | 7.62 | 1.41 |
| Missing | 0.65 | 0.4346 | 46.02 | 7.52 | 1.39 |
| 82 | 0.63 | 0.4346 | 48.40 | 7.40 | 1.37 |
| 43 | 0.65 | 0.4346 | 42.21 | 7.04 | 1.30 |
| 64 | 0.63 | 0.4346 | 44.20 | 6.83 | 1.26 |
| 109 | 0.63 | 0.4346 | 44.20 | 6.81 | 1.26 |
| 83 | 0.61 | 0.4346 | 45.90 | 6.74 | 1.24 |
| Missing | 0.61 | 0.4346 | 44.20 | 6.45 | 1.19 |
| 88 | 0.59 | 0.4346 | 44.20 | 5.99 | 1.11 |
| 61 | 0.57 | 0.4346 | 42.67 | 5.41 | 1.00 |
| 46 | 0.52 | 0.4346 | 42.67 | 4.48 | 0.83 |
| 30 | 0.52 | 0.4346 | 42.67 | 4.42 | 0.82 |
| 67 | 0.49 | 0.4346 | 42.67 | 4.06 | 0.75 |
| 69 | 0.49 | 0.4346 | 42.67 | 4.04 | 0.75 |
| Missing | 0.49 | 0.4346 | 42.67 | 4.00 | 0.74 |
| 112 | 0.48 | 0.4346 | 42.67 | 3.90 | 0.72 |
| 33 | 0.47 | 0.4346 | 42.67 | 3.63 | 0.67 |
| 110 | 0.46 | 0.4346 | 42.67 | 3.60 | 0.66 |
| 60 | 0.45 | 0.4346 | 42.67 | 3.43 | 0.63 |
| Missing | 0.45 | 0.4346 | 42.67 | 3.35 | 0.62 |
| 75 | 0.41 | 0.4346 | 42.67 | 2.85 | 0.53 |
| **Average** | **0.75** |  | **45.09** | **10.47** | **1.93** |
